# Investigating the Propagation of 0.1 - 2.5 THz Radiation Through a Phantom Ear Model: Implications for Wireless Network-Biological Tissue Interaction

**DOI:** 10.1101/2023.08.15.552876

**Authors:** Reza Shams, David Sly, Zoltan Vilagosh

## Abstract

This research focuses on the investigation of the propagation of frequencies between 0.1 and 2.5 THz through a phantom ear model using terahertz (THz) time-domain spectroscopy (TDS). While the use of THz frequencies between 0.1 to 0.3 THz in fifth and sixth generation cellular networks has gained significant attention, there is also a growing interest in utilising higher frequencies, such as 1 THz and above, for various applications, including the Internet of Things (IoT), autonomous vehicles, smart sensors, and smart cities. Despite the limited absorption coefficient of soft tissues at 5G and 6G frequencies (0.2-0.4 mm), the effect of higher frequencies on deeper regions of the ear, such as the tympanic membrane (with a thickness of 0.1 mm), has not been extensively studied. The study aims to determine the optimal conditions for THz transmission through the ear canal and to investigate the interaction between wireless networks and biological tissues. The results show that when parallel to the ear canal, the average power flux density within the central region of the tympanic membrane is 97% of the incident excitation. However, the outer ear structures are highly protective, with less than 0.4% of the power flux density directed towards them reaching the same region. Due to the sensitivity of the tympanic membrane to mechanical changes, in-vivo assessments are necessary to evaluate the penetration of THz frequencies into the ear canal, assess the suitability of current radiation safety limits, and evaluate the implications of devices that emit these frequencies. The study highlights the importance of understanding the interaction between THz radiation and biological tissues, particularly in the context of emerging wireless technologies, and the need for further research to ensure their safety and effectiveness.

## Introduction

As the implementation of 5G technology in sub-terahertz frequencies (30 to 90 GHz) progresses, and the development of 6G mobile networks is underway to utilize even higher frequencies up to 0.3 THz, the potential applications of this new spectrum are expanding (Samsung, 2020; Yang et al., 2019). Areas such as autonomous vehicles, robotics, and the Internet of Things (IoT) are benefitting from this technology to connect sensors, equipment, and processing units in smart homes and cities. However, the deployment of these applications may increase human exposure to these frequencies, necessitating a thorough understanding of their potential effects on human health and safety, and appropriate risk mitigation measures (Bandara et al., 2020).

Water’s absorption of terahertz (THz) and sub-terahertz frequencies is well-established and plays a crucial role in the behavior of these frequencies in various applications, including telecommunications, sensing, and spectroscopy. Biological tissues, such as the skin, cornea, and tympanic membrane, with high water content, experience strong absorption of incident radiation within the first few millimeters of tissue. Consequently, THz waves primarily interact with the surface layers of these tissues. The penetration depth of THz radiation, around 0.4 THz, has been found to be approximately 0.3 mm (You et al., 2018), which may not be sufficient to effectively halt the propagation of higher THz frequencies through structures such as the tympanic membrane.

As the frequency of electromagnetic waves increases, the ability of the ear canal and pinna to prevent their propagation into the inner ear decreases. Theoretical predictions suggest that very high frequencies have a higher chance of penetrating the ear canal and interacting with sensitive structures within the ear, such as the tympanic membrane. Vilagosh et al. (2019) modelling suggests that the penetration of sub-terahertz frequencies within the human ear can vary depending on the incident angle, with the power flux density within the tympanic membrane region decreasing as the incident angle deviates from the normal. This is particularly relevant in real-world scenarios, as THz radiation can propagate from various angles.

Additionally, the penetration depth of THz radiation decreases with increased frequency, with a maximum penetration depth of around 0.3 mm at 0.4 THz. This means that as the frequency increases, the radiation is less likely to penetrate deep into the tissue and will primarily interact with the surface layers. In the case of the tympanic membrane, with a thickness of only 0.1 mm, it is theoretically possible for high frequency radiation to be attenuated by this structure. However, the effect of THz frequencies on the deeper parts of the ear, such as the tympanic membrane, has not been extensively studied and empirical assessments are needed to fully understand the interaction of these frequencies with the ear canal.

The use of sub-terahertz frequencies in 5G technology and the potential for even higher frequencies in 6G networks may impact human exposure due to the strong absorption of incident radiation by living soft tissues. The tympanic membrane may be a significant point of interaction for THz radiation, as indicated by Vilagosh et al. (2019) computer modeling research. Therefore, it is crucial to conduct further studies to confirm these findings and determine the potential implications of exposure to THz radiation, particularly regarding incident angle and the relationship between frequency and penetration depth.

Therefore, in this study, two hypotheses were proposed. The first hypothesis stated that the highest amount of THz radiation would be detected at the surface of the tympanic membrane when the radiation was incident parallel to the ear canal. The second hypothesis postulated that the transmission of THz radiation through the tympanic membrane would decrease with an increase in frequency. To evaluate these hypotheses, an ear model was employed to investigate the effect of THz radiation incident at various angles on the ear. Measurements were taken of the resulting radiation at the tympanic membrane surface. Additionally, the transmission of THz radiation beyond the tympanic membrane was examined to determine absorption and penetration at different frequencies.

## Method

To replicate the water content of a human ear canal, an ear model was constructed using paper and soaked in distilled water for approximately two minutes. The dimensions of the model were based on an average human ear, and the water content was verified through weight measurements before and after soaking. Industrial weigh scales (A&D FX-300i) with a sensitivity of 0.001g were employed to ensure an accurate representation of the water content within the model. This approach facilitated the replication of the ear canal conditions and provided a suitable sample for examining THz radiation transmission.

**Figure 1.**
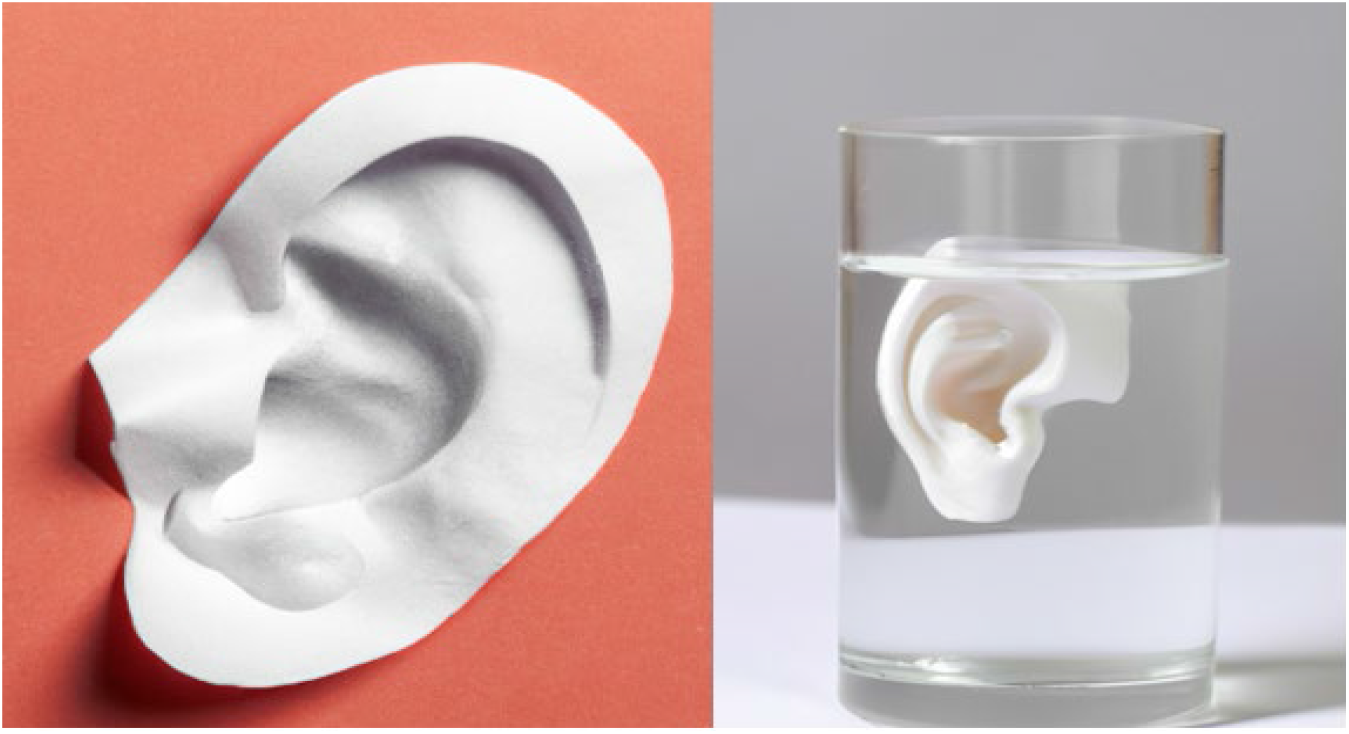
Ear Model Made of Paper Soaked in Water. *Note:* Left figure shows the dry ear model, and the right figure shows the soaked model in distilled water.

A thin layer of paper was employed as a tympanic membrane model. As water is known to be the primary interactor with THz radiation, the paper was soaked to achieve a water composition of 80%, which closely resembles the water consistency of a natural tympanic membrane. This approach enabled the accurate simulation of THz radiation absorption and penetration through the tympanic membrane in a controlled experimental setting.

The BATOP THz-TDS system was utilised to generate THz radiation with a frequency range of 0.1 to 2.5 THz. The accompanying software provided data on the amplitude of the radiation detected at various frequencies, facilitating a comprehensive analysis of the radiation transmission and absorption by the ear model. The system was operated under controlled conditions to ensure the accuracy and reliability of the results.

Gold mirrors were employed as wave guides to adjust the angle of incident radiation in this study. As depicted in the figure, altering the mirror position enabled the modification of the incident radiation angle relative to the ear. The reflective properties of gold at THz frequencies were considered, as gold exhibits high reflectivity in the THz range, making it an ideal material for this experiment. The gold mirrors were meticulously chosen and aligned to ensure precise control of the THz radiation’s incident angle.

**Figure 2.**
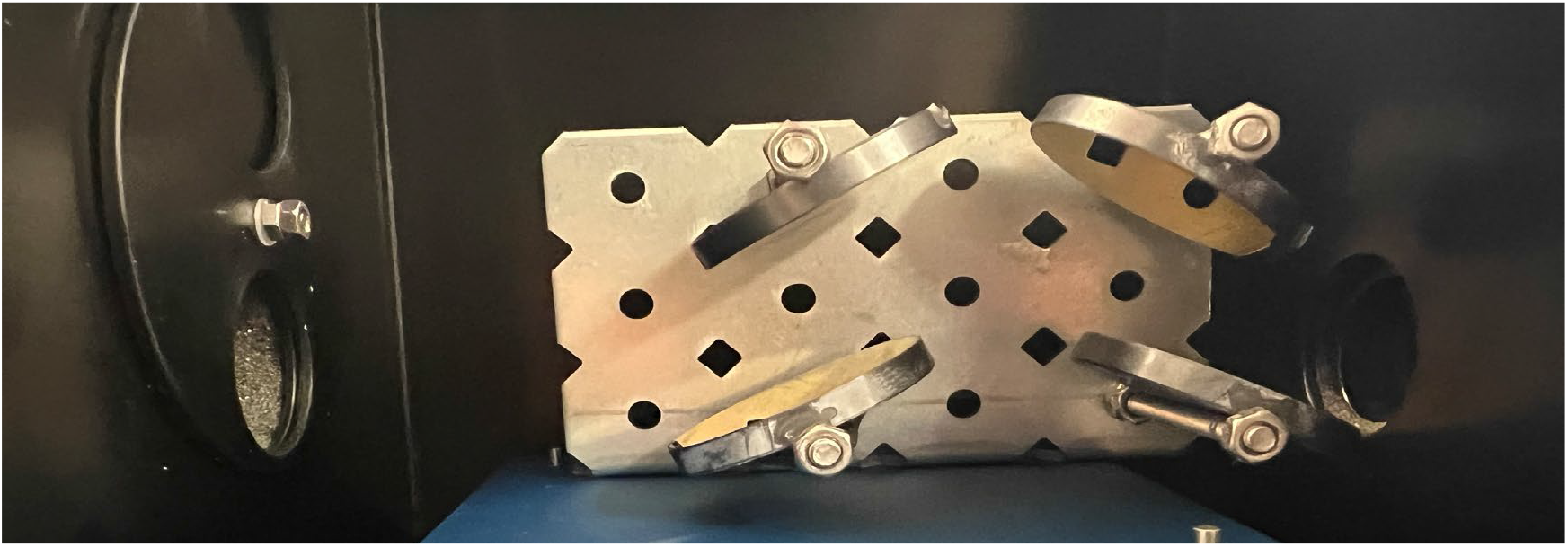
Wave Guide Made of Four Gold Mirrors to Change the Angle of Incident. *Note:* The mirrors used in the experiment were positioned at a 45-degree angle to effectively reflect the THz radiation. The last mirror, located at the bottom left, was adjustable to allow for changes in the angle of reflection.

The ear model, created using paper and soaked in water to achieve a water composition similar to that of the human ear canal, was then weighed to ensure the correct water-to-paper ratio. A background measurement with the sample was acquired before positioning the ear model within the BATOP THz-TDS system, which was used to generate THz radiation with a frequency range of 0.1 to 2.5 THz. To change the angle of incident radiation, four gold mirrors were employed as wave guides. The mirror was first positioned at an angle to reflect THz parallel to the ear canal, followed by positioning the mirror at angles of 30, 45, and 60 degrees relative to the ear canal. The gold mirrors were selected due to their reflective properties at THz frequencies. Data on the amplitude detected at various frequencies was recorded using the included software.

During the data analysis stage, the Fast Fourier Transform (FFT) of the signal at various frequencies was obtained through the use of T3DS software. Subsequently, the transmission through the ear model was calculated using the Beer-Lambert Law. This involved determining the ratio of the incident wave amplitude in the presence of the sample to the incident wave amplitude in the absence of the sample. This methodology allowed for the quantification of the absorption and penetration of THz radiation through the ear model.

## Results

The study aimed to investigate the absorption and transmission of THz radiation through the ear and evaluate two hypotheses concerning the impact of THz radiation on the tympanic membrane. The ear model employed in the study was constructed using paper and soaked in distilled water to replicate the water content of the human ear canal. As illustrated in Figure 3, the water content of the paper model played a significant role in the absorption of THz radiation. The soaked paper model exhibited a high absorption coefficient for THz radiation, consistent with the high-water content of living soft tissues.

**Figure 3.**
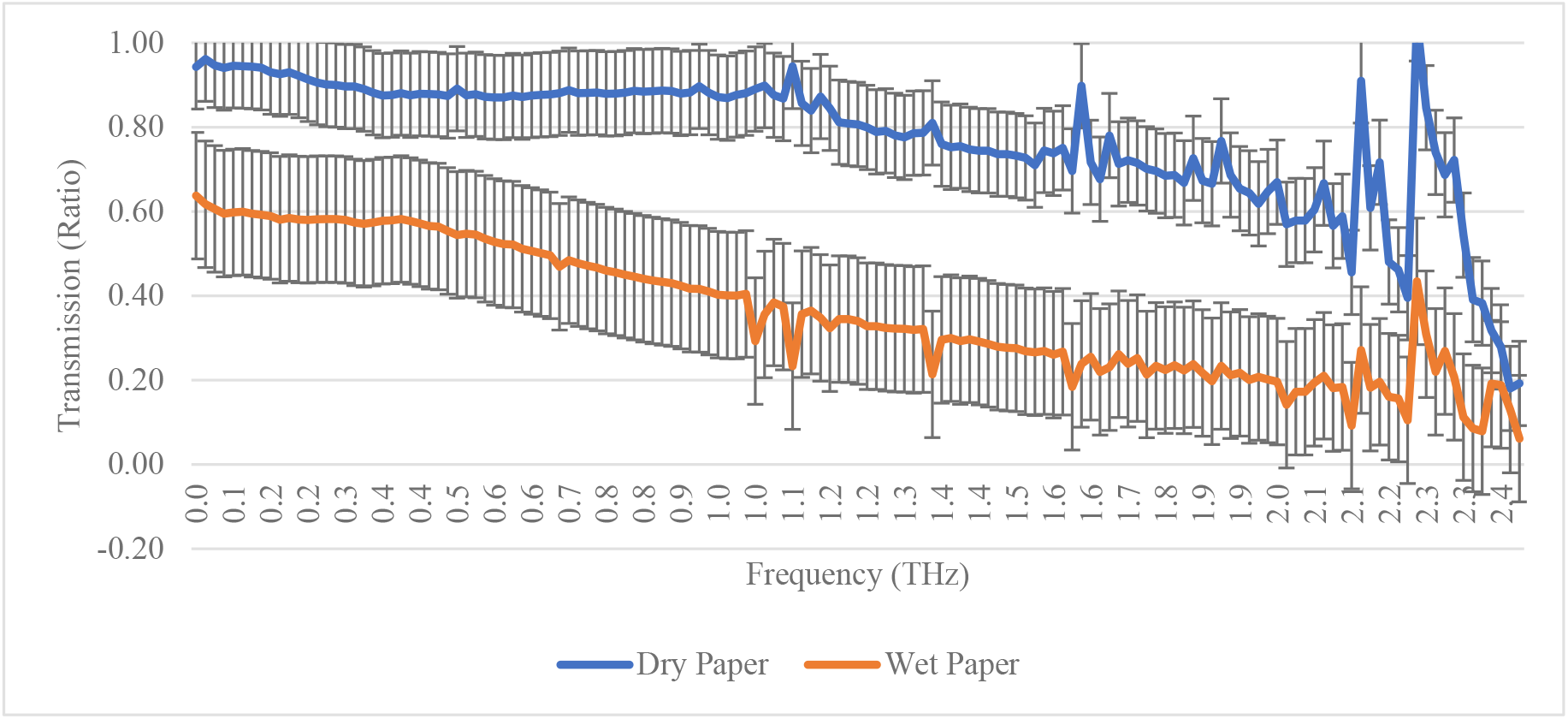
Dry and Soaked Paper THz Transmission Spectra. *Note:* As shown dry paper shows no attenuation at lower frequencies but increases at higher frequencies, however soaked paper shows significantly lower transmission, suggesting that the bulk of the radiation is either absorbed or reflected.

**Figure 4.**
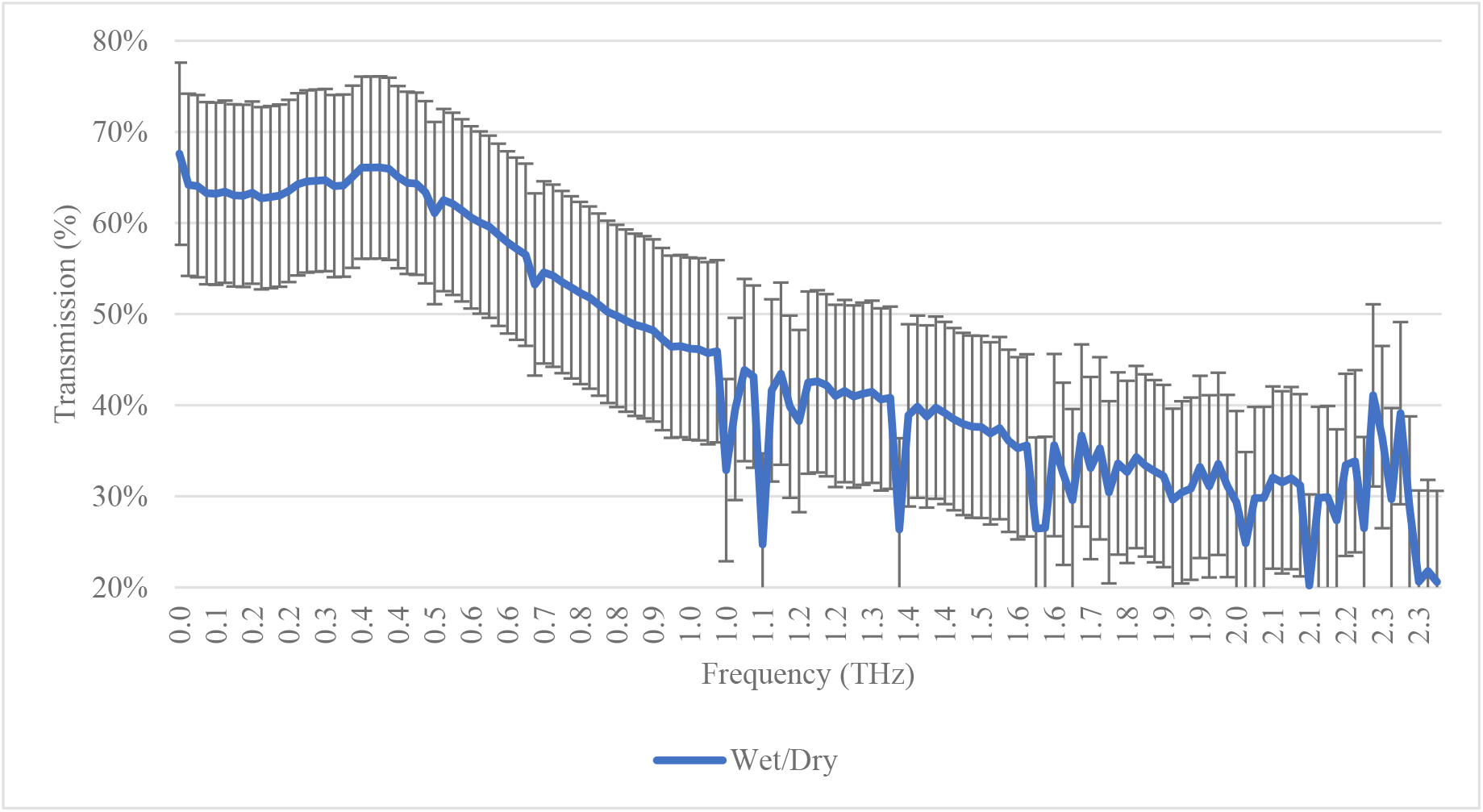
Normalised Transmission (%) of Soaked Paper / Dry Paper. *Note:* Transmission decreases with frequency, at 0.1 THz 68% of the radiation is transmitted, while at higher frequency such as 2 THz only 30% of the total radiation is transmitted and the rest absorbed or reflected.

Transmission properties of paper vary according to its moisture content and the radiation frequency. Dry paper demonstrates minimal to no attenuation at lower frequencies, but experiences increased attenuation at higher frequencies. Conversely, soaked paper displays considerably lower transmission, indicating that a substantial portion of the radiation is either absorbed or reflected by the paper. Moreover, the behaviour of soaked paper in response to THz radiation varies with frequency, with lower THz frequencies generally exhibiting higher transmission percentages than higher frequencies. At lower frequencies, THz radiation is transmitted with reduced efficiency, whereas at higher frequencies, the transmission decreases, implying that THz radiation is either absorbed or reflected by the material.

Furthermore, the soaked paper as an alternative model for studying the interaction of THz radiation with human skin demonstrates that the model exhibits characteristics akin to human skin, with comparable levels of absorption and transmission of THz radiation..

The angle of incidence of the THz radiation significantly influenced the amount of radiation detected on the other side of the ear phantom model. Parallel excitation to the ear canal provided the highest incident radiation detected on the opposite side of the model. As shown in Figure 5, a greater percentage of incident radiation reached the tympanic membrane when the incident angle was parallel to the ear canal, with the peak occurring at around 1 THz but continuing to decline at higher frequencies.

**Figure 5.**
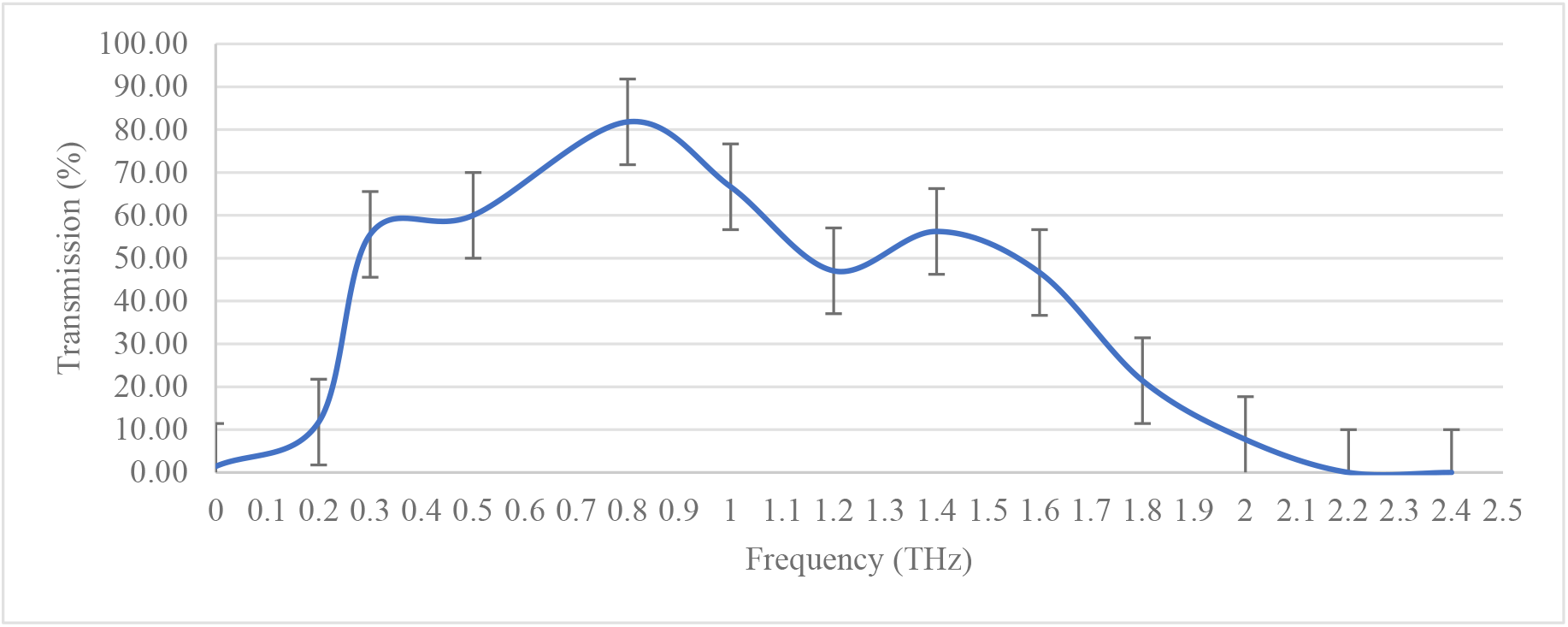
Normalised Transmission (%) of 0.1 to 2.5 THz at Parallel to the Ear Canal. *Note:* Transmission at lower frequencies is very low but increases with higher frequencies.

Moreover, results at a 30-degree angle revealed that a smaller proportion of the incident wave could propagate through the ear canal. As illustrated in Figure 6, radiation transmission decreased as the angle of incidence deviated from the parallel orientation to the ear canal. At a 45-degree angle, THz radiation transmission was negligible, remaining closer to 0%

**Figure 6.**
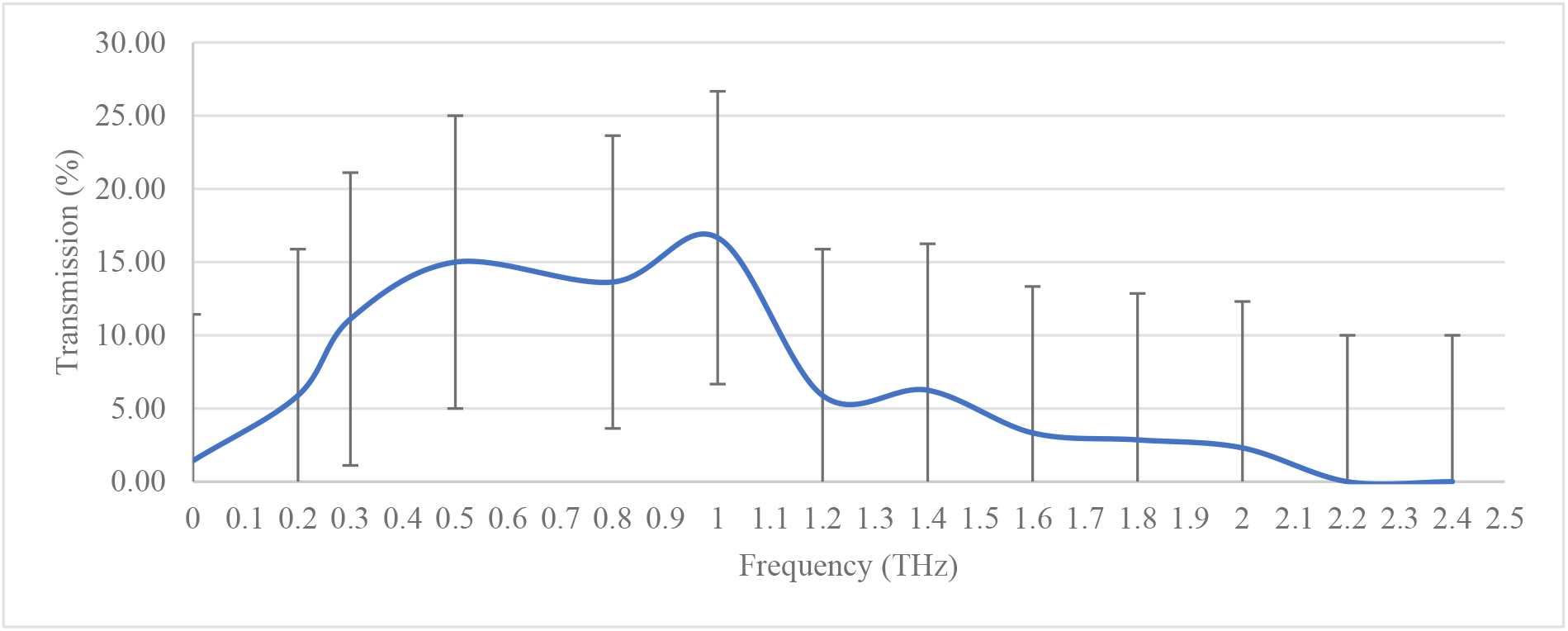
Normalised Transmission (%) of 0.1 to 2.5 THz at 30 Degrees Angle to the Ear Canal. *Note:* Transmission at lower frequencies is very low but increases with higher frequencies. transmission.

## Discussion

The results of this study highlight the need to consider the potential impact of THz radiation on human exposure and health, especially as these frequencies are increasingly utilised in telecommunications. Living soft tissues, such as skin and tympanic membrane, have a high water content, which results in strong absorption of incident THz radiation, primarily through absorption. The absorption coefficient is significant, increasing from 75 cm^-1^ at 0.1 THz to 100 cm^-1^ at 0.3 THz. The study revealed that the parallel excitation to the ear canal provided the highest incident radiation detected on the other side of the ear phantom model, which may have implications for the directionality of THz radiation in real-world scenarios. Consequently, it is essential to conduct further research to comprehend the potential health risks related to exposure to these frequencies, particularly concerning the incident angle and the relationship between frequency and penetration depth.

Utilising paper soaked in water as an alternative model for studying the interaction of THz radiation with human skin demonstrated similar characteristics to human skin, with comparable levels of reflection and transmission of THz radiation. The soaked paper model allowed for the adjustment of water content to examine the interaction more conveniently than using real ear tissue. Employing this model marks a significant advancement in understanding THz radiation interaction with human skin, enabling a more accurate assessment of potential health risks associated with exposure to these frequencies.

The angle of incidence of THz radiation significantly influenced the amount of radiation detected on the other side of the ear phantom model. As exposure to THz radiation can occur at any angle, understanding the parameters, such as the incident angle, is vital to minimize tympanic exposure. This study’s findings indicate that the penetration of sub-terahertz frequencies within the human ear varies depending on the incident angle, with parallel excitation resulting in the highest propagation. The ear canal and pinna, which comprise the outer structures of the ear, act as barriers to prevent the propagation of lower frequencies into the inner ear. Nevertheless, as the frequency of electromagnetic waves increases, the ability of the ear canal and pinna to prevent propagation diminishes. This is attributable to higher frequencies having a shorter wavelength, enabling them to penetrate through smaller openings and gaps. Therefore, comprehending the impact of THz radiation on the human ear and the potential health risks related to exposure to these frequencies is crucial, particularly as the use of sub-terahertz frequencies in 5G technology and the potential for even higher frequencies in 6G networks continue to expand.

This study offers insights into the absorption and transmission of THz radiation through the ear and its potential implications for human exposure and health. The parallel excitation to the ear canal provided the highest incident radiation detected on the other side of the ear phantom model, which may have implications for the directionality of THz radiation in real-world scenarios. The use of soaked paper as an alternative model for studying the interaction of THz radiation with human skin represents a significant advancement, allowing for a more precise assessment of potential health risks related to exposure to these frequencies. The angle of incidence of THz radiation had a substantial impact on the amount of radiation detected on the other side of the ear phantom model, emphasizing the importance of understanding the angle of incidence in real-world scenarios. Further research is necessary to determine the potential health risks associated with exposure to THz radiation, particularly regarding the incident angle and the relationship between frequency and penetration depth.

## Refrence

Bandara, P., Chandler, T., Kelly, R., McCredden, J., May, M., Weller, S., … & Wojcik, D. (2020). 5G wireless deployment and health risks: Time for a medical discussion in Australia and New Zealand. Journal of the Australasian College of Nutritional and Environmental Medicine, 39(2), 27–34.

Samsung. (2020). Samsung’s 6G White Paper Lays Out the Company’s Vision for the Next Generation of Communications Technology. https://news.samsung.com/global/samsungs-6g-white-paper-lays-out-the-companys-vision-for-the-next-generation-of-communications-technology

Vilagosh, Z., Lajevardipour, A., & Wood, A. (2019). Computational simulations of the penetration of 0.30 THz radiation into the human ear. Biomed Opt Express, 10(3), 1462–1468. https://doi.org/10.1364/BOE.10.001462

Vilagosh, Z., Lajevardipour, A., & Wood, A. (2019). An empirical formula for temperature adjustment of complex permittivity of human skin in the terahertz frequencies. Bioelectromagnetics, 40(1), 74–79. https://doi.org/10.1002/bem.22156

Vilagosh, Z., Lajevardipour, A., & Wood, A. W. (2019). Computational phantom study of frozen melanoma imaging at 0.45 terahertz. Bioelectromagnetics, 40(2), 118–127. https://doi.org/10.1002/bem.22169

Yang, P., Xiao, Y., Xiao, M., & Li, S. (2019). 6G wireless communications: Vision and potential techniques. IEEE Network, 33(4), 70–75. https://doi.org/10.1109/MNET.2019.1800418

You, B., Chen, C. Y., Yu, C. P., Wang, P. H., & Lu, J. Y. (2018). Frequency-dependent skin penetration depth of terahertz radiation determined by water sorption-desorption. Opt Express, 26(18), 22709–22721. https://doi.org/10.1364/oe.26.022709

